# Functional insights into *Plasmodium* actin depolymerizing factor interactions with phosphoinositides

**DOI:** 10.1101/2024.11.29.626011

**Authors:** Devaki Lasiwa, Inari Kursula

**Author notes:** Corresponding authors: Devaki Lasiwa; Inari Kursula.

## Abstract

Malaria is caused by protozoan parasites, *Plasmodium* spp., that belong to the phylum Apicomplexa. The life cycle of these parasites depends on two different hosts; the definitive host, or vector, is a mosquito, and the intermediate host is a vertebrate, such as human. Malaria parasites use a unique form of substrate-dependent motility for host cell invasion and egress, which is dependent on an actomyosin motor complex called the glideosome. Apicomplexa have a small set of actin regulators, which are poorly conserved compared to their equivalents in higher eukaryotes. Actin depolymerizing factors (ADFs) are key regulators responsible for accelerating actin turnover in eukaryotic cells. The activity of ADFs is regulated by membrane phosphoinositides. Malaria parasites express two ADF isoforms at different life stages. ADF1 differs substantially from canonical ADF/cofilins and from *Plasmodium* ADF2 in terms of both structure and function. Here, we studied the interaction of both *Plasmodium* ADFs with phosphoinositides using biochemical and biophysical methods and mapped their binding sites on ADF1. Both *Plasmodium* ADFs bind to different phosphoinositides, and binding *in vitro* requires the formation of vesicles or micelles. Interaction with phosphoinositides increases the α-helical content of the parasite ADFs, and the affinities are in the micromolar range. The binding site for PI(4,5)P2 in *Pf*ADF1 involves a small, positively charged surface patch.

## Introduction

ADF/cofilins are among the most central proteins that regulate cell proliferation, migration, polarity, and dynamic regulation of organ morphology (1, 2). ADF/cofilins control actin dynamics by accelerating actin polymerization and depolymerization *via* their severing activity as well as by nucleation (3–5). Upon bnding to ADP-G-actin, ADF/cofilin proteins inhibit nucleotide exchange to ATP-G-actin (6). ADF/cofilins are regulated by a plethora of mechanisms, including phosphorylation/dephosphorylation, pH, and interactions with other proteins and phosphoinositides. Phosphorylation of a conserved serine residue at the N-terminus of ADF/cofilins inhibits their binding to F-actin (7–9). The severing and depolymerization properties are regulated by pH, and ADF/cofilins typically are most active at high pH (10–12).

Phosphatidylinositol and its phosphorylated derivatives, phosphoinositides, are multifunctional lipids involved in the modulation of many cellular events, such as signal transduction, regulation of membrane traffic, cytoskeleton, and the permeability and transport functions of membranes (13, 14). The inositol ring of phosphatidylinositols can be reversibly phosphorylated at positions 3, 4, and 5, which results in the formation of seven possible phosphoinositide species. phosphatidylinositol 4,5-bisphosphate [PI(4,5)P_2_] affects the actin cytoskeleton by interacting directly with several actin-binding proteins. PI(4,5)P_2_ binding inhibits proteins involved in actin filament disassembly and activates nucleation or polymerization promoting proteins (15–18). The interaction of ADF/cofilins with phosphoinositides is a regulatory mechanism that mainly occurs at the plasma membrane. ADF/cofilins are among the few actin-binding proteins present at the leading edge of migrating cells, highlighting the importance of ADF/cofilin-membrane interactions. Both PI(4,5)P_2_ and phosphadylinositol 3,4,5-triphosphate [PI(3,4,5)P_3_] interact with ADF/cofilins with a relatively high affinity compared to other phosphoinositides and have been suggested to act as phospoinositide density sensors (19, 20). The binding of phosphosinositides to ADF/cofilins occurs by electrostatic interactions through a large, positively-charged surface area and is mutually exclusive with actin binding (19, 21, 22).

Apicomplexa, a diverse group of largely obligate intracellular parasites, including *Plasmodium* spp., comprise significant pathogens of animals, including humans. They display a unique form of substrate-dependent motility to cross non-permissive biological barriers (migration), invade the target host (invasion), and exit from infected cells (egress). All these depend on an actomyosin motor (22, 23). Apicomplexan parasites have a limited number of actin-binding proteins controlling actin filament turnover (23, 24).

Unlike other apicomplexan parasites, *Plasmodium* spp. have two ADF isoforms: ADF1 and ADF2. ADF1 is expressed during all lifecycle stages and implicated in cell motility (25, 26). ADF2, on the other hand, is expressed in the sexual stages of the parasite lifecycle (27). Both ADF1 and ADF2 share the core fold with canonical ADF/cofilins. However, there are marked differences between the *Plasmodium* ADFs compared to each other and to canonical ADF/cofilins. The largest difference in the *Plasmodium* ADFs appears in the C-terminal half, which is involved in interactions with G- and F-actin. A hairpin loop (called F-loop) connecting β-strands 4 and 5 is shorter in ADF1 than in ADF2 and in other ADF/cofilins. β-strand 6 connecting α-helix 3 to the C-terminal helix is missing in ADF1, and has a shorter α-helix 4. In addition, hydrophobic residues implicated in G-actin binding are missing in ADF1 (26, 28). All in all, ADF2 is more similar to canonical ADF/cofilins than ADF1.

Several phosphoinositides have been detected in *P. falciparum* infected erythrocytes such as phosphoinositide 3-phosphate (PI3P), phosphoinositide 4-phosphate (PI4P), PI(4,5)P_2_, and low levels of phosphoinositide 3,4-bisphosphate [PI(3,4)P_2_] and PI(3,4,5)P_3_ (29). A previous study using recombinant *Pf*ADF1 has shown a low micromolar affinity for phosphatidylinositol derivatives (26). However, little is known about how *Plasmodium* ADFs interact with phosphoinositides and their specificities. Here, we used biochemical and biophysical techniques to study the interactions of *Plasmodium* ADFs with different phosphoinositides.

## Results

### *Plasmodium* ADFs bind phosphoinositides specifically

The interaction of *Plasmodium* ADFs with different phosphoinositides was assessed by incorporating 10% phosphoinositides into 1-palmitoyl-2-oleoyl-sn-glycero-3- phosphatidylcholine (POPC) vesicles. Co-sedimentation assays for both *Plasmodium* ADFs showed hardly any binding to POPC vesicles without phosphoinositides, but in the presence of phosphoinositides, the binding was significantly enhanced (Figure 1A and 1B). 18-25% of *Pf*ADF1 sedimented in the presence of all different phosphoinositides (Figure 1A and 1C), suggesting either a lack of specificity or a limitation of this method. For *Pb*ADF2, the lowest binding was observed for PI(4,5)P_2_ (Figure 1D); only approximately 10% of *Pb*ADF2 sedimented with PI(4,5)P_2_, compared to 13-28% with the other phosphoinositides (Figure 1B and 1D).

**Figure 1:**
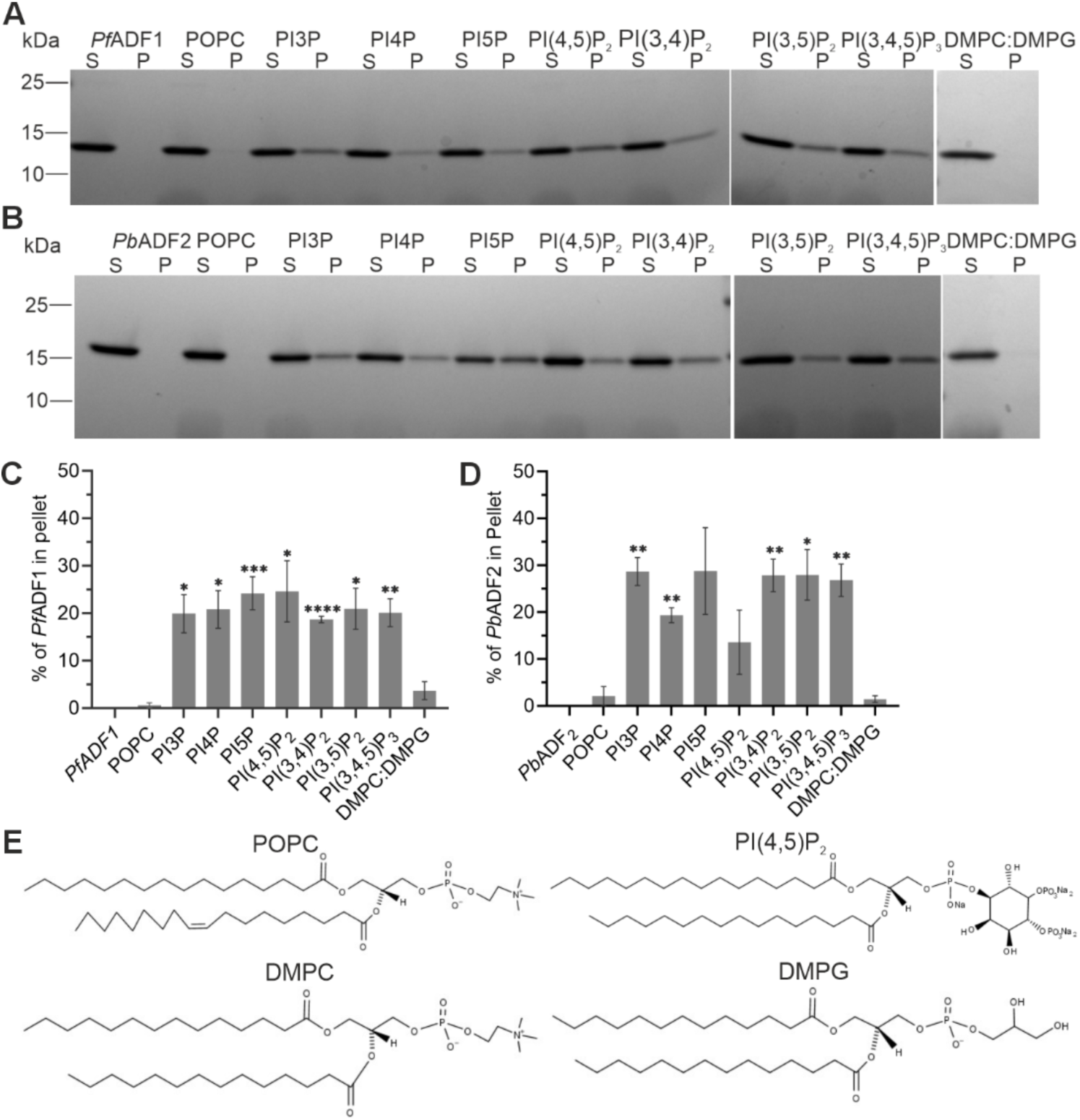
*Plasmodium* ADFs interact with different phosphoinositides. A representative SDS-PAGE analysis of the vesicle co-sedimentation assay of *Pf*ADF1 (A) and *Pb*ADF2 (B), with various 10% phosphoinositides, POPC, and DMPC:DMPG vesicles. S and P denote the supernatant and pellet, respectively. Bar diagrams showing the proportion of *Pf*ADF1 (C) and *Pb*ADF2 (D) in the pellet fractions (n = 3) quantitated from figures 1A and 1B, respectively. Data were plotted as mean ± standard error mean (SEM). For both, C and D, asterisks represent statistical significances determined with unpaired two-tailed t-tests against the POPC control. \**P* < 0.05, \*\**P* < 0.01, \*\*\**P* < 0.001, \*\*\*\**P* < 0.0001. (E) Molecular structures of POPC, PI(4,5)P_2_, DMPC, and DMPG.

As phosphoinositides have negatively charged head groups (Figure 1E), it was investigated, whether ADFs also bind to other negatively charged vesicles. For this purpose, negatively charged 1,2-dimyristoyl-sn-glycero-3phosphocholine (DMPC):1,2-dimyristoyl-sn-glycero-3-phospho-glycerol (DMPG) (1:1) (Figure 1E) vesicles were used. Interestingly, neither of the *Plasmodium* ADFs bound to DMPC:DMPG vesicles more than to POPC (Figure 1C and 1D), suggesting that binding of the *Plasmodium* ADFs to phosphoinositides is specific to the head groups.

### Phosphoinositide binding induces conformational changes in *Plasmodium* ADFs

We used synchrotron radiation circular dichroism (SRCD) spectroscopy to study whether the conformation of *Plasmodium* ADFs is affected by interaction with phosphoinositides. All phosphoinositide vesicles, except PI(3,4,5)P_3_ and PI(3,4)P_2,_ increased the α-helical content of *Pf*ADF1, while the POPC control vesicles did not, as indicated by an increase in the positive absorption peak at 195 nm and the two negative absorption peaks at 208 nm and 222 nm (Figure 2 A and 2B, Table 1). The largest changes occurred at 195 nm. Deconvolution of the SRCD spectra between 180 and 250 nm using the beta structure selection (BeStSel) server (30) showed an increase of α-helix content up to 1.5-fold seen upon interaction with PI3P (Table 1). All phosphoinositides, except PI(3,4)P_2_, increased the α-helical content of *Pb*ADF2, PI(3,5)P_2_ having the most prominent effect. To investigate whether the micellar structure of the phosphoinositides in solution is important for the ADF binding, the effect of the soluble form SPI(4,5)P_2_ *i.e* dibutanoyl phosphatidylinositol 4,5-bisphosphate, on the secondary structure of the *Pf*ADF1 was also tested. SPI(4,5)P_2_ did not affect the protein conformation (Figure 2B). In addition, we did not detect any interaction between SPI(4,5)P_2_ and *Pf*ADF1 using isothermal titration calorimetry. Thus, the vesicle environment seems to be required for the binding of *Pf*ADF1 to phosphoinositides.

**Figure 2:**
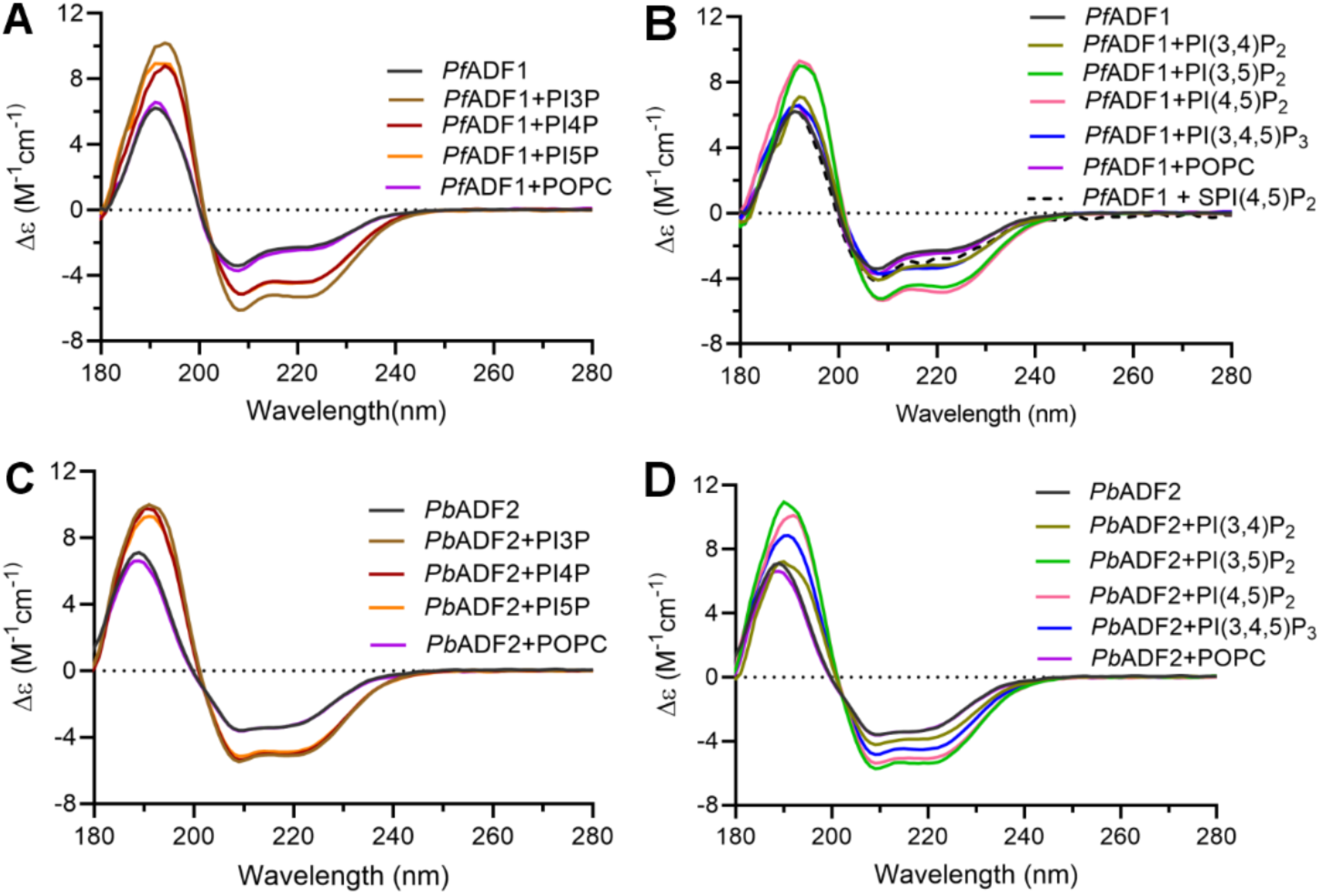
SRCD spectra of *Plasmodium* ADFs in the presence of various phosphoinositides. SRCD spectra of *Pf*ADF1 with and without different (A) monophosphoinositide vesicles and (B) di- and triphosphoinositide vesicles. SRCD spectra of *Pb*ADF2 with and without different (C) monophosphoinositide vesicles, and (D) di- and triphosphoinositide vesicles.

**Table 1:** Deconvolution of SRCD spectra of *Plasmodium* ADFs with different phosphoinositides.

| Protein ± phosphoinositide vesicles | Secondary structure content (%) | | | |
| --- | --- | --- | --- | --- |
| | α-helix | β-strand | turn | other |
| *Pf*ADF1 | 24 | 22 | 11 | 43 |
| *Pf*ADF1 + POPC | 24 | 20 | 11 | 46 |
| *Pf*ADF1 + PI3P | 41 | 13 | 7 | 39 |
| *Pf*ADF1 + PI4P | 38 | 12 | 10 | 40 |
| *Pf*ADF1 + PI5P | 40 | 8 | 10 | 42 |
| *Pf*ADF1 + PI(3,4)P2 | 27 | 21 | 12 | 40 |
| *Pf*ADF1 + PI(3,5)P2 | 36 | 17 | 7 | 40 |
| *Pf*ADF1 + PI(4,5)P2 | 39 | 13 | 10 | 38 |
| *Pf*ADF1 + PI(3,4,5)P3 | 27 | 22 | 10 | 41 |
| *Pf*ADF1 + SPI(4,5)P2 | 22 | 21 | 12 | 46 |
| *Pf*ADF1 (PDB ID: 2XF1) | 32 | 26 | 5 | 38 |
| *Pb*ADF2 | 19 | 36 | 10 | 36 |
| *Pb*ADF2 + POPC | 16 | 36 | 10 | 38 |
| *Pb*ADF2+ PI3P | 42 | 17 | 3 | 38 |
| *Pb*ADF2+ PI4P | 37 | 18 | 5 | 40 |
| *Pb*ADF2+ PI5P | 36 | 19 | 5 | 40 |
| *Pb*ADF2+ PI(3,4)P2 | 29 | 28 | 5 | 38 |
| *Pb*ADF2+ PI(3,5)P2 | 41 | 16 | 4 | 39 |
| *Pb*ADF2+ PI(4,5)P2 | 40 | 15 | 7 | 38 |
| *Pb*ADF2 + PI(3,4,5)P3 | 31 | 25 | 6 | 38 |
| *Pb*ADF2 (PDB ID: 2XFA) | 31 | 27 | 9 | 33 |

### Both *Pf*ADF1 and *Pb*ADF2 bind to phosphoinositides with a micromolar affinity

Both *Plasmodium* ADFs contain single tryptophan residues (Trp-26 in *Pf*ADF1 and Trp-92 in *Pb*ADF2). The Trp-92 in *Pb*ADF2 is at positions similar to tryptophans seen in conventional ADF/cofilins (Figure 6). Thus, intrinsic tryptophan fluorescence spectroscopy was used to determine the binding affinity of the *Plasmodium* ADFs to different phosphoinositides (Figure 3). The background fluorescence from the phosphoinositide vesicles alone was monitored by phosphoinositide vesicle titrations. These showed an increased fluorescence intensity but no shift of the maxima (Figure S1). The fluorescence intensity of these vesicle titrations was subtracted from the protein phosphoinositide titration, and the changes in tryptophan fluorescence *versus* phosphoinositide concentration were plotted.

**Figure 3:**
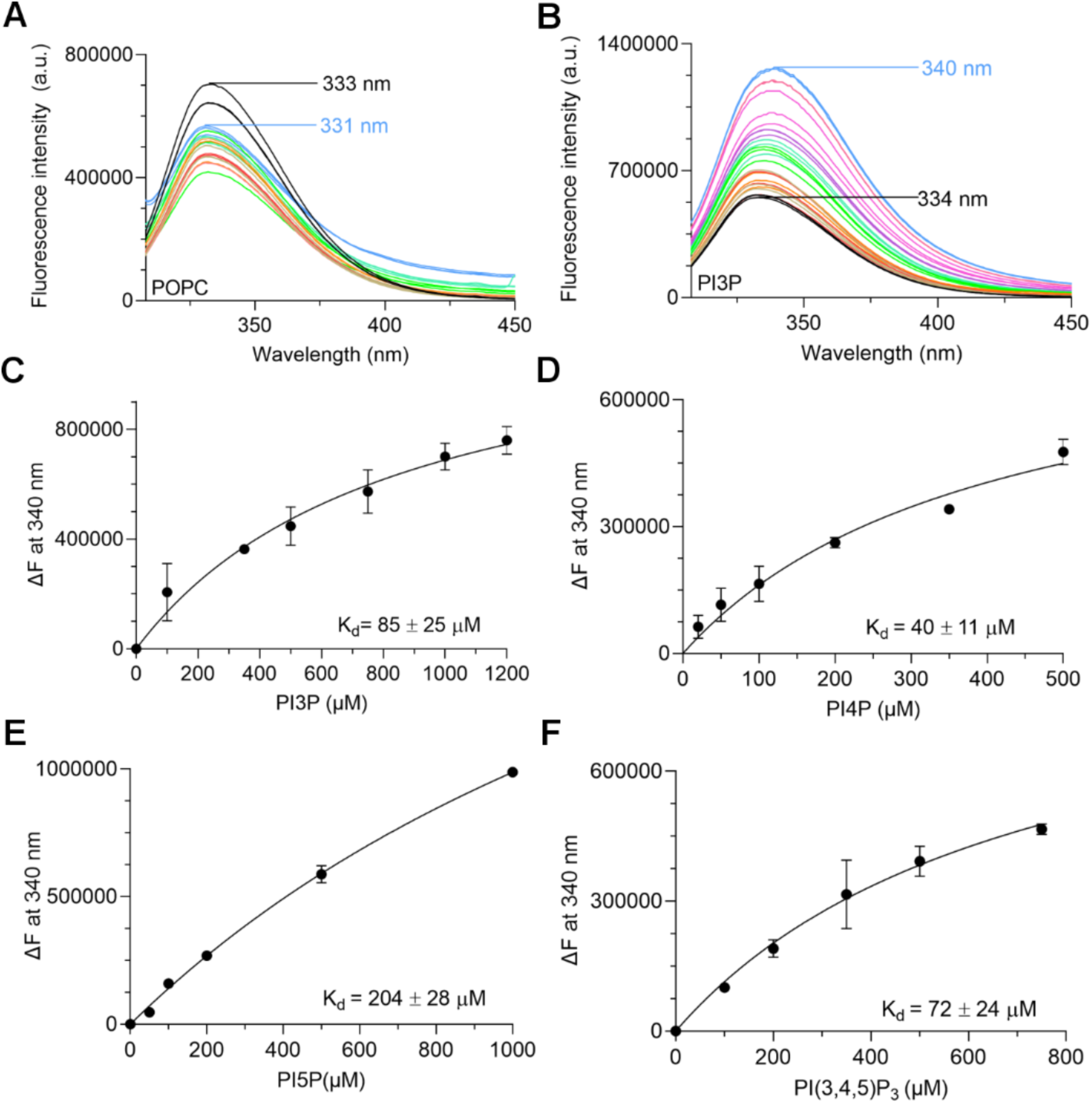
Determination of *Pf*ADF1-mono/triphosphoinositide interactions using tryptophan fluorescence. *Pf*ADF1 fluorescence intensity in the presence of increasing concentrations of POPC (A), and PI3P (B) vesicle were recorded for the first (no POPC/phosphoinositide, black) and last (blue) titration curves. The emission of *Pf*ADF1 decreased upon increasing POPC concentration but increased upon increasing phosphoinositide concentrations. The phosphoinositide concentration *vs.* change in fluorescence intensity at the maximum emission wavelength were plotted and analyzed using a one-site specific binding model to obtain the binding affinities for PI3P (C), PI4P (D), PI5P (E), and PI(3,4,5)P_3_ (F). Data were plotted as mean ± SEM from three independent experiments.

When aqueous solutions of *Pf*ADF1 were excited at 295 nm, emission spectra with maxima at 334 nm were obtained (Figure 3A). The titration of *Pf*ADF1 with phosphoinositides resulted in 5-7 nm shifts of the fluorescence emission maxima to longer wavelengths (red shift) in a concentration-dependent manner. A red shift typically results from a tryptophan becoming more exposed to solvent. In addition, titration of *Pf*ADF1 with phosphoinositides also increased the peak intensity. A POPC vesicle titration showed a reduction of tryptophan fluorescence intensity without any shifts of the maxima (Figure 3A), and it was not possible to determine a dissociation constant (K_d_) for the interaction between *Pf*ADF1 and POPC, suggesting no binding or a very weak interaction. All the phosphoinositides had K_d_s in the range of 40 to 200 µM. Of the different phosphoinositide vesicles, PI(3,5)P_2_ and PI4P had the highest affinities and PI5P had the lowest affinity (Figure 3 and 4).

**Figure 4:**
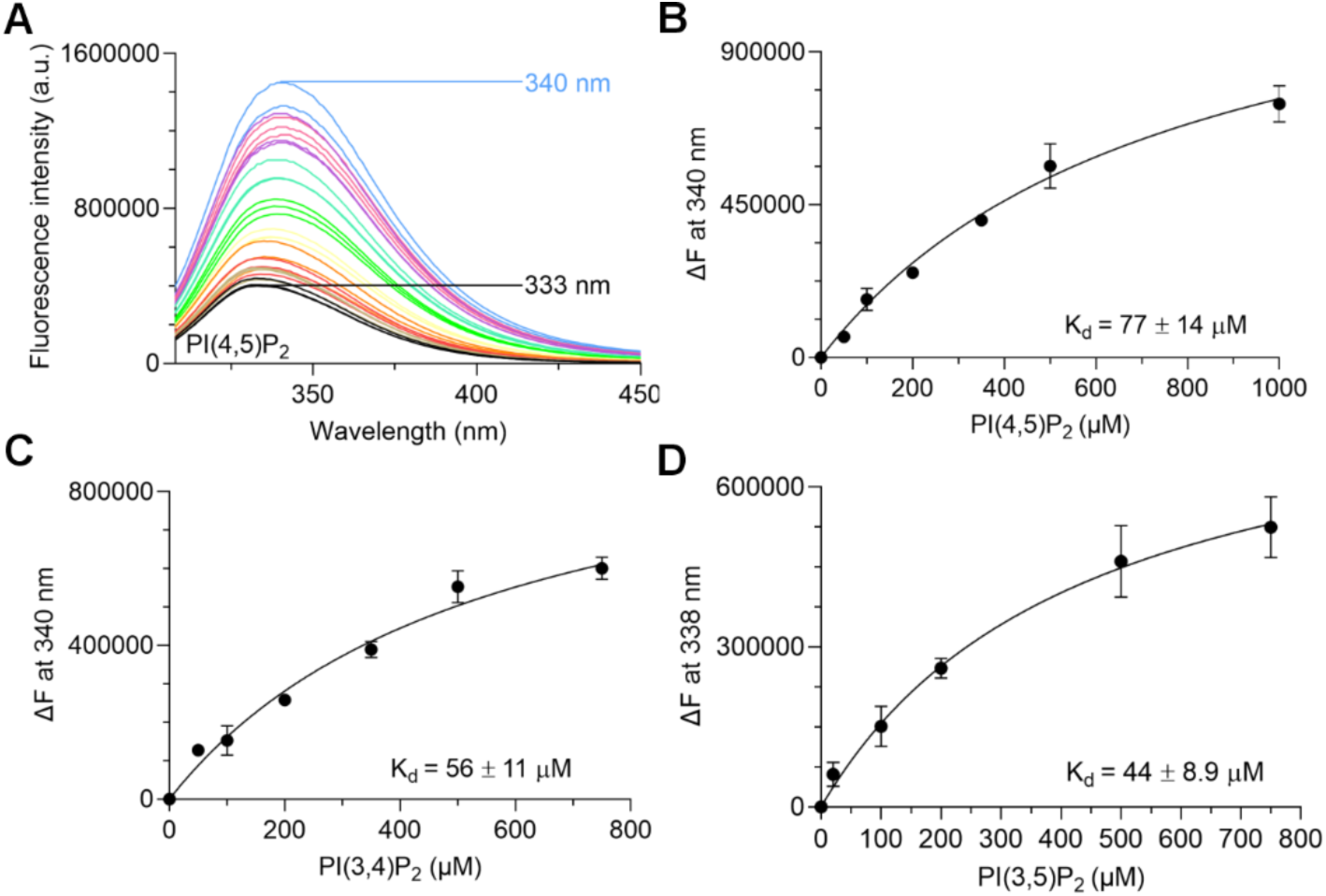
Determination of *Pf*ADF1-diphosphoinositide interactions using tryptophan fluorescence. *Pf*ADF1 fluorescence intensities in the presence of increasing concentrations of PI(4,5)P_2_ (A) were recorded for the first (no phosphoinositide, black) and last (blue) titration curves. The emission of *Pf*ADF1 increased upon increasing phosphoinositide concentration. The phosphoinositide concentration *vs.* change in fluorescence intensity at the maximum emission wavelength were plotted and analyzed using a one-site specific binding model to obtain the binding affinities for PI(4,5)P_2_ (B), PI(3,4)P_2_ (C), and PI(3,5)P_2_. Data were plotted as mean ± SEM from three independent experiments.

When *Pb*ADF2 was excited at 295 nm, emission spectra with maxima at 328 nm were obtained (Figure 5A). The titration of *Pb*ADF2 with phosphoinositides resulted in red shifts of around 6 nm and a concentration-dependent increase in fluorescence intensity (Figure 5B). In contrast to *Pf*ADF1, *Pb*ADF2 titration with POPC vesicles alone resulted in an increase in fluorescence intensity in concentration-dependent manner but no shifts of the maxima were observed, and the binding affinity for POPC could not be determined. The K_d_ of *Pb*ADF2 binding to PI(4,5)P_2_ vesicles was 12 ± 5.6 µM (Figure 5D), which indicates a higher affinity than observed for *Pf*ADF1 to any of the phosphoinositide vesicles (Figure 3 and 4). We could not determine the binding affinity for PI5P vesicles, although we observed a similar shift in the fluorescence maximum as seen for PI(4,5)P_2_, suggesting weak binding (Figure 5C).

**Figure 5:**
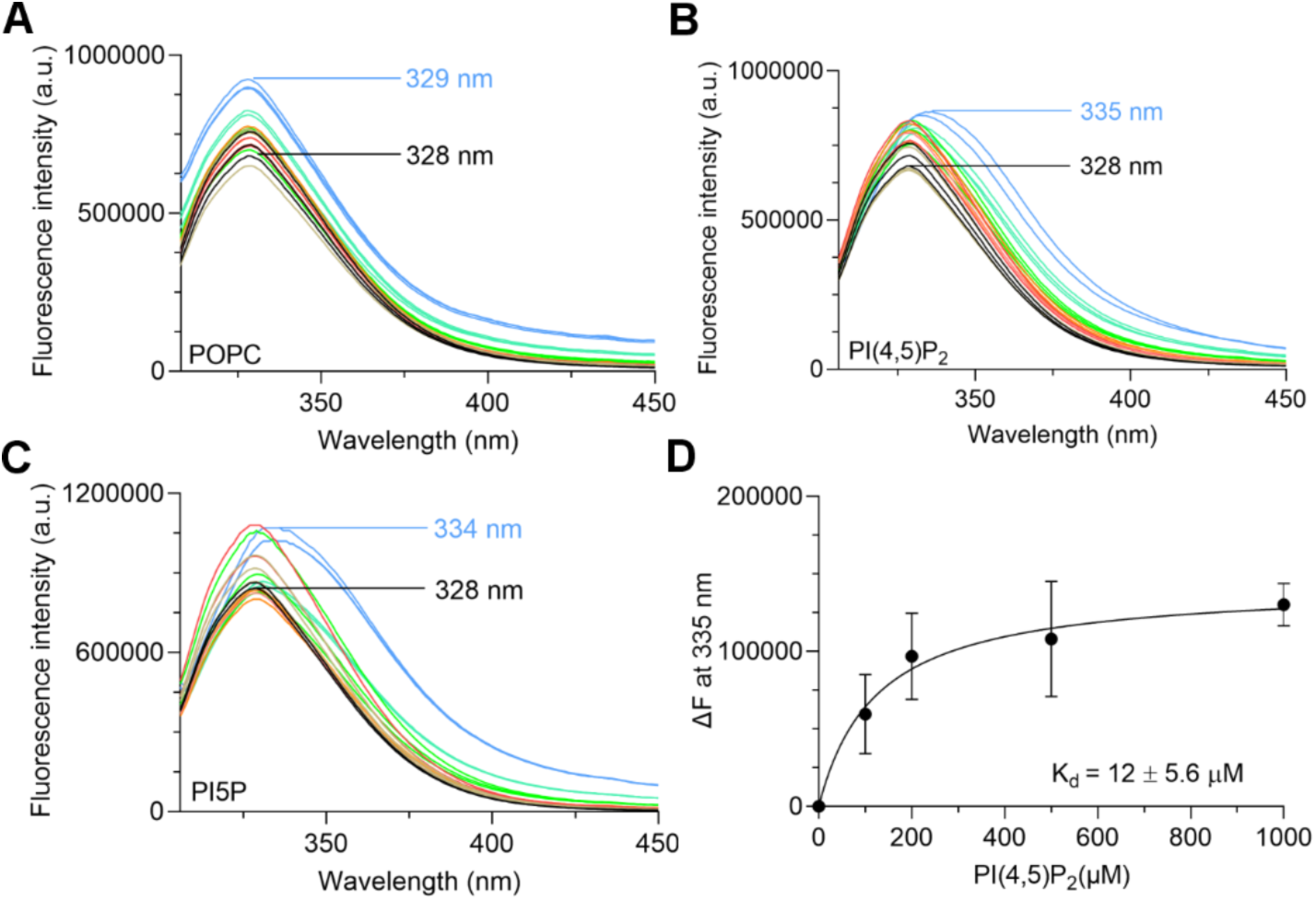
Determination of *Pb*ADF2-phosphoinositide interactions using tryptophan fluorescence. *Pb*ADF2 fluorescence intensities in the presence of increasing concentrations of POPC (A), PI(4,5)P_2_ (B), and PI5P (C) vesicle were recorded for the first (no POPC/phosphoinositide, black) and last (blue) titration curves. The emission of *Pb*ADF2 increased upon increasing POPC and phosphoinositide concentrations. The phosphoinositide concentration *vs.* change in fluorescence intensity at the maximum emission wavelength of 340 nm were plotted and analyzed using a one-site specific binding model to obtain the binding affinity for PI(4,5)P_2_ (D). Data were plotted as means ± SEM from three independent experiments.

### Mapping the PI(4,5)P_2_ binding site on *Pf*ADF1

Previous studies on chicken and yeast cofilins have been conducted using NMR and native gel electrophoresis to map the phosphoinositide-binding sites. These techniques have limitations, as NMR chemical shift experiments were carried out using water-soluble di-C8 forms of PI(4,5)P_2_, and native gel electrophoresis assays were performed under nonphysiological conditions (20, 31). Thus, we combined mutagenesis of *Pf*ADF1 with CD and fluorometric assays to shed light on the phosphoinositide binding mode. The previous results suggested that phosphoinositide binding with *Pf*ADF1 requires formation of vesicles or micelles, which would be most consistent with binding *via* electrostatic interactions. Positively charged residues in four clusters important for phosphoinositide binding in other ADF/cofilins were mutated to alanine or glutamine, guided by a multiple sequence alignment (Figure 6). These mutations are located on the surface of *Pf*ADF1 (Figure 7C). In addition, we mutated Ser-3, a phosphorylation target, to either glutamate to mimic phosphorylation at this site, or an alanine to remove the polar side chain. All mutants were properly folded, as indicated by SRCD (Figure S2).

**Figure 6:**
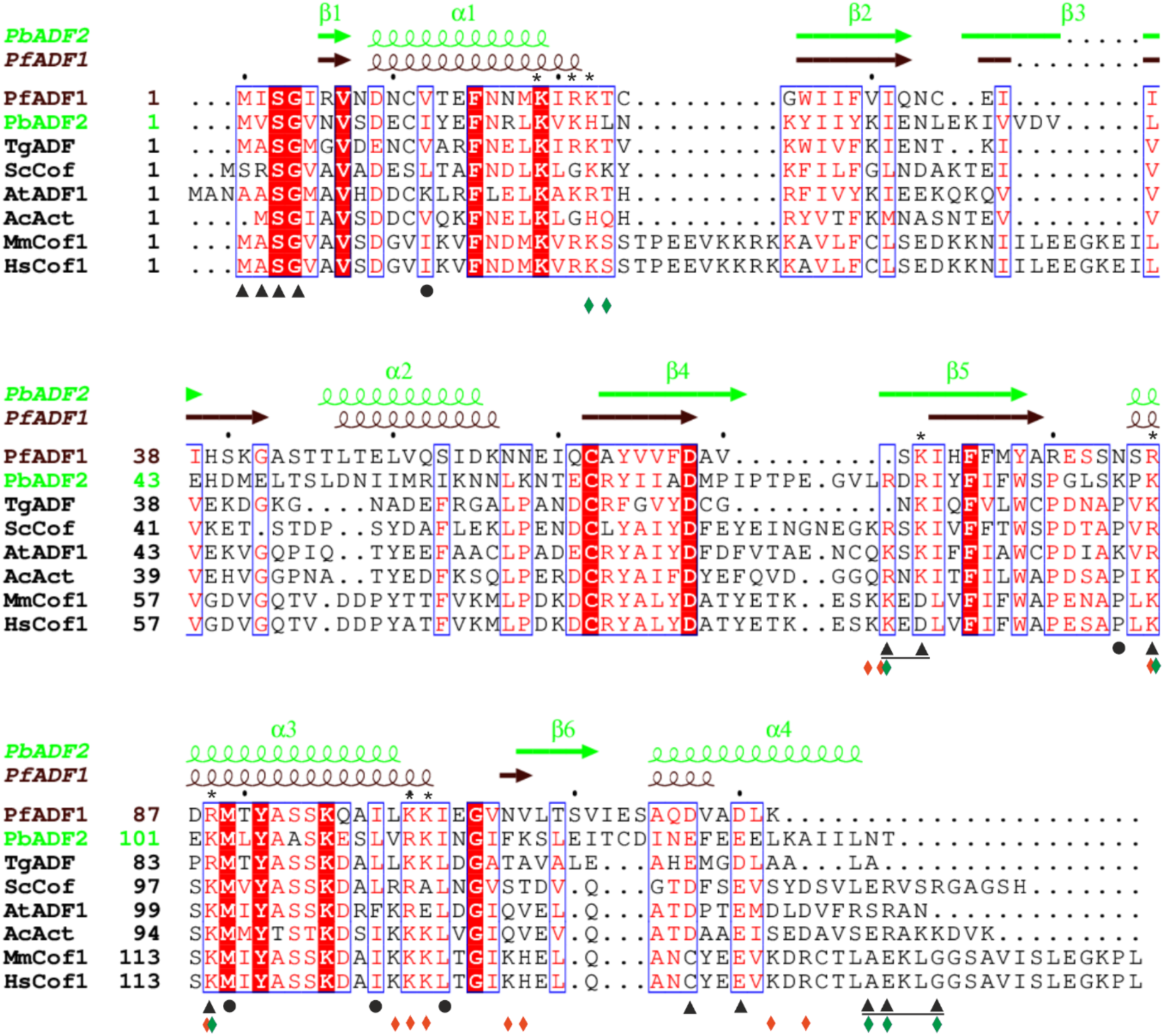
Effect of mutations on *Pf*ADF1 binding to PI(4,5)P2 determined using tryptophan fluorescence. (A) K72A mutant fluorescence intensity in the presence of increasing concentrations of PI(4,5)P_2_ were recorded for the first (no phosphoinositide, black) and last (blue) titration curves. The emission of K72A increased upon increasing phosphoinositide concentration. (B) The phosphoinositide concentration *vs.* change in fluorescence intensity of *Pf*ADF1 and its mutants at the maximum emission wavelength of 340 nm were plotted and analyzed using a one-site specific binding model to obtain the binding affinity for PI(4,5)P2. The data were plotted as mean ± SEM from three independent experiments. (C) Electrostatic surface potential of *Pf*ADF1 [PDB ID: 2XFA,(28)]. The amino acids mutated in this study are indicated. Shown are two different orientations 90° apart (red negative, blue positive, and white, neutral).

**Figure 7:**
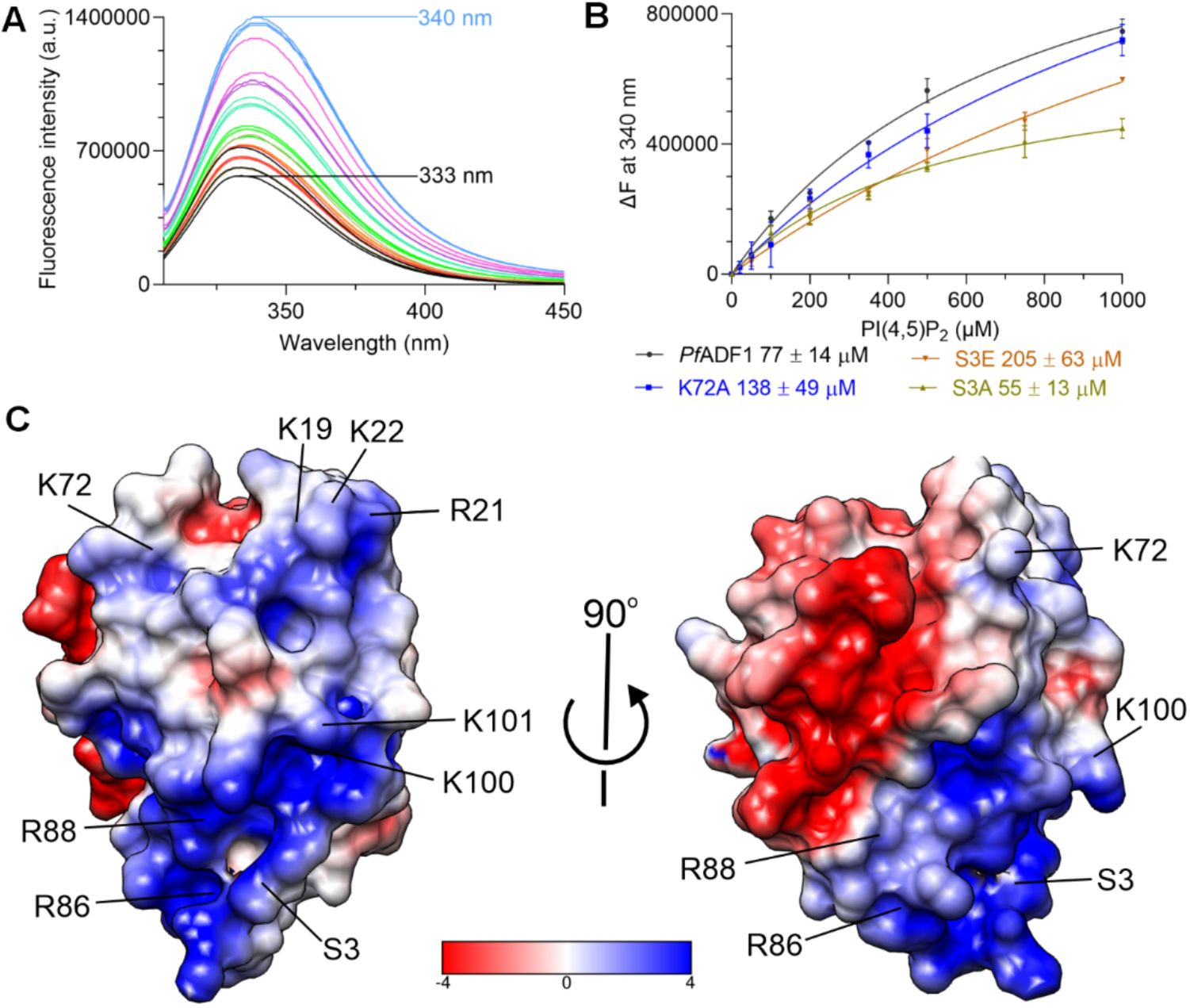
Multiple sequence alignment of *Plasmodium* ADFs and selected other ADF/cofilins. The amino acid sequences of *Plasmodium* ADFs were aligned with other ADF/cofilin family members using ClustalW2 (52). Strictly conserved residues are shown in red boxes, and regions of residues with similar properties are indicated with blue boxes. The secondary structure elements of *Pf*ADF1 and *Pb*ADF2 are shown above the alignment in black and green, respectively. G-actin-binding sites identified in yeast cofilin by mutagenesis (20) and synchrotron footprinting (54) are marked with black triangles and circles, respectively. Residues involved in the F-actin-binding site are marked with underlined black triangles. The phosphoinositide-binding sites identified by mutagenesis in yeast cofilin (34) and mouse cofilin-1 (55) are marked with green and red diamonds, respectively. The *Pf*ADF1 residues mutated in this study are indicated by asterisks above the sequences. The sequences include those of *P. falciparum* ADF1 (*Pf*ADF1), *P. berghei* ADF2 (*Pb*ADF2), *T. gondii* ADF (*Tg*ADF), *S. cerevisiae* cofilin (*Sc*Cof), *A. thaliana* ADF1 (*At*ADF1), *A. castellanii* actophorin (*Ac*Act), *M. musculus* cofilin-1 (MmCof), and *H. sapiens* cofilin (*Hs*Cof).

The effect of the mutations on phosphoinositide binding was studied using PI(4,5)P_2_. Based on the SRCD results, *Pf*ADF1 mutations did not have notable effects on binding of PI(4,5)P_2_ (Figure S2). However, in the tryptophan fluorescence assay (Figure 7B), residues Lys-19, Arg-21, and Lys-22 and Arg-86 and Arg-88 of α3 helix seemed to be critical for PI(4,5)P_2_ binding, as binding was abolished in the alanine mutants. The same was seen for the Lys-100 and Lys-101 mutations to alanine and glutamine (Table 2). This suggests that these residues may contribute to the PI(4,5)P_2_ binding. The S3A mutation did not affect the phosphoinositide-binding. However, substitution of serine with glutamate decreased the binding to PI(4,5)P_2_ (Table 2), indicating that phosphorylation might negatively affect phosphoinositide binding. Overall, these results suggest that the PI(4,5)P_2_ binding site on *Pf*ADF1 may be a small positively charged patch located concentrated around these surface residues (Figure 7C).

**Table 2:**
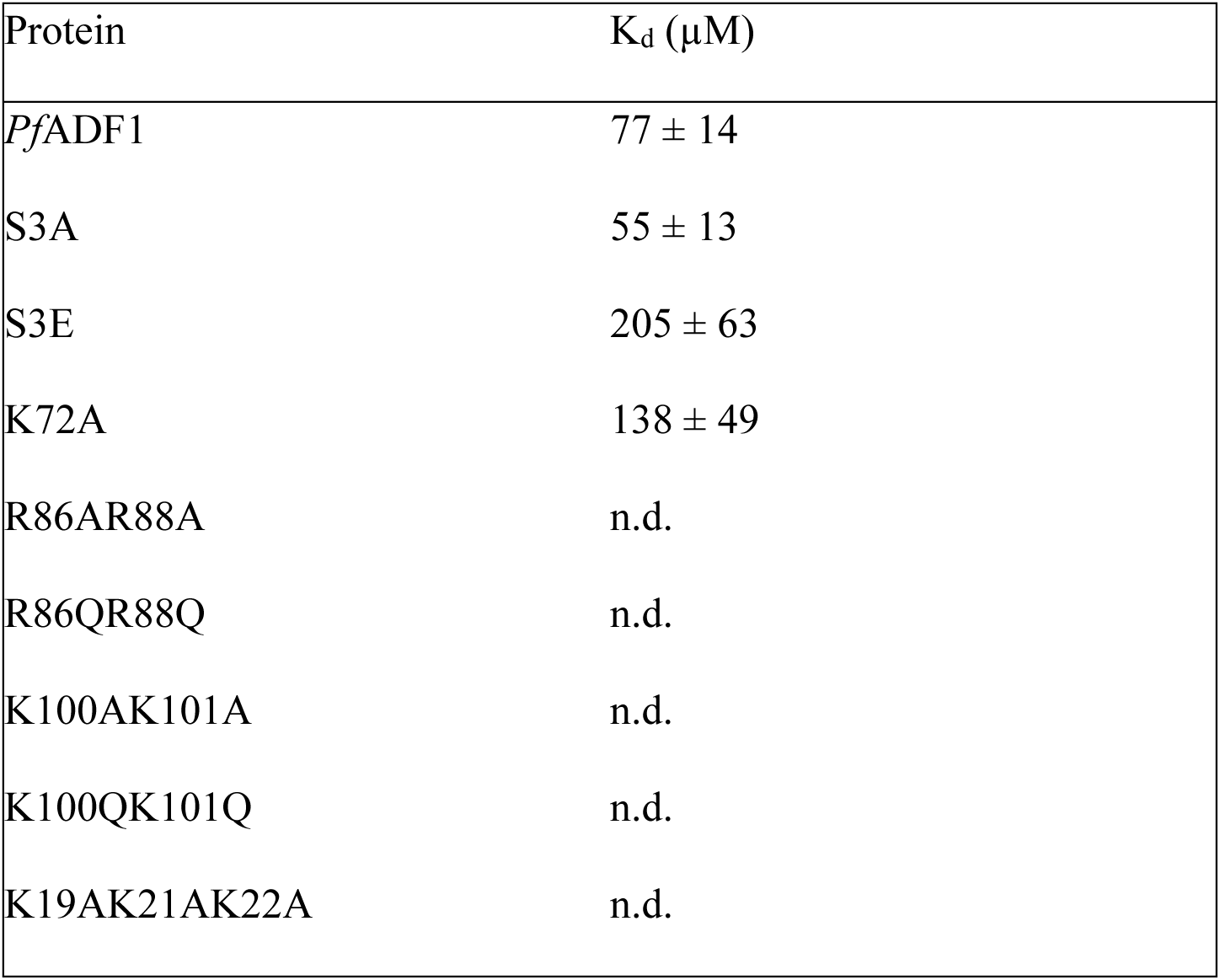
Binding affinities of *Pf*ADF1 and several mutants to PI(4,5)P2 determined using the tryptophan fluorescence assay. Data are plotted as mean ± SEM (n = 3) (n.d. = not determined).

## Discussion

Unlike other Apicomplexa, *Plasmodium* spp. express two ADFs: ADF1 and ADF2. They are present at different life stages of the parasite, indicating different functions. While ADF1 is essential for parasite viability and invasion and is present throughout the parasite lifecycle, ADF2 is not expressed during the erythrocytic stages (25). Although phosphoinositides only account for 1% of total cellular lipids in eukaryotic cells, they are critical in various cellular processes, such as signal transduction, cell motility, cytoskeletal organization and membrane transport. They are not present uniformly in cell membranes but enriched in certain cellular compartments (32). ADF/cofilins are regulated by phosphoinositides *via* inhibition of actin-binding activity because the phosphoinositide and actin-binding sites partially overlap. ADF/cofilins typically bind multiple PI(4,5)P_2_ head groups through a large, positively charged protein interface (19). In the case of *Plasmodium*, the regulation of the ADFs by phosphoinositides has not been studied before.

Here, we studied whether and how the *Plasmodium* ADFs bind to different phosphoinositides. PI(4,5)P_2_ is the most abundant phosphoinositide in the plasma membrane in human and likely to play a dominant regulatory role over other phosphoinositides in the organization of the actin cytoskeleton (32). However, other phosphoinosides also interact with ADF/cofilins, and little is known about membrane compositions in the apicomplexan membranes and during different life cycle stages. Both *Plasmodium* ADFs bind different phosphoinositides in POPC vesicles (Figure 2). Interestingly, water-soluble short-chain phosphoinositides did not affect the conformation of the *Plasmodium* ADFs, indicating that the membrane environment or micelle/vesicle formation is important for binding. The curvature of the membrane and ratio of phosphoinositide to other lipids have been reported to be important for phosphoinositide binding in other cytoskeletal proteins (33, 34). These results are consistent with previous findings in the *Toxoplasma gondii* ADF (*Tg*ADF), which did not show any interaction with dioctanoyl PI(4,5)P_2_ in the ITC or NMR experiments (35). This might be due to the lack of formation of vesicles or micelles, as a water-soluble phosphoinositide was used.

A previous study on yeast and chicken cofilins showed that they have stronger interactions with di- and tri-phosphorylated phosphoinositides than the monophosphorylated forms (19, 31). In contrast, we could not show a clear preference among the various phosphoinositides in the lipid co-sedimentation or tryptophan fluorescence assays, but CD spectroscopy showed that monophosphorylated phosphoinositides induced a larger conformational change upon binding to the ADFs, than di- and tri-phosphorylated phosphoinositide (Figure 2, Table 1). This was particularly the case for PI3P. This may have *in vivo* relevance during parasite infection. The production of phosphoinositides increases after infection of red blood cells with *P. falciparum*, particularly for PI3P, PI4P, and PI(4,5)P_2_. Of the total PI-monophosphates in infected erythrocytes, 30% are PI3P. PI3P levels do not fluctuate in most eukaryotic cells, but in *P. falciparum*, there is an approximately four-fold increase in the proportion of them to total phosphoinositides. It plays a role in hemoglobin uptake, biogenesis of the apicoplast, and artemisinin resistance. Normally, unicellular organisms do not produce PI(3,4,5)P_3_, however, the presence of PI(3,4,5)P_3_ in the *P. falciparum* schizonts, and the *P. berghei* ookinetes has been confirmed (36). The affinity of PI5P to *Pf*ADF1 was 204 µM, while it could not be determined for *Pb*ADF2, suggesting no binding or a very weak interaction. In line with this, PI5P has been found only in asexual blood stages of malarial parasites (37), where ADF2 is not present. PI(4,5)P_2_ is involved in calcium signaling cascades as a substrate for phospholipase C in sporozoite gliding motility, merozoite egress, and male gametocyte exflagellation (38). An increase in the α-helical content upon binding to phosphoinositides has been previously observed for profilin and gelsolin. CD studies of gelsolin synthetic peptides showed that they undergo coil-to-helix transition upon PI(4,5)P_2_ binding (39, 40).

A number of *Plasmodium* ADF1 mutants, in which a charged residues were mutated to alanine, or glutamine, were constructed to map the PI(4,5)P_2_ binding site on the *Pf*ADF1. Three of the mutants (K19AR21AK22A, R86AR88A, and K100AK101A) did not strongly interact with PI(4,5)P_2_. These sites reside in α1 and the long α3 helix of *Pf*ADF1, respectively, suggesting that the *Pf*ADF1 might not have a defined phosphoinositide-binding pocket. Therefore, the *Pf*ADF1 might simultaneously interact with more than one phosphoinositide molecule or does not contain a specific interaction site. In support of this, the positioning of proteins in membrane (PPM) Webserver (41) predicted that N-terminus and α3 helix of both *Pf*ADF1 and *Pb*ADF2 interact with membranes (Figure 8). Similar observations have been reported for mouse cofilin-1, yeast cofilin, and human cofilin-1 (19, 20, 42).

**Figure 8:**
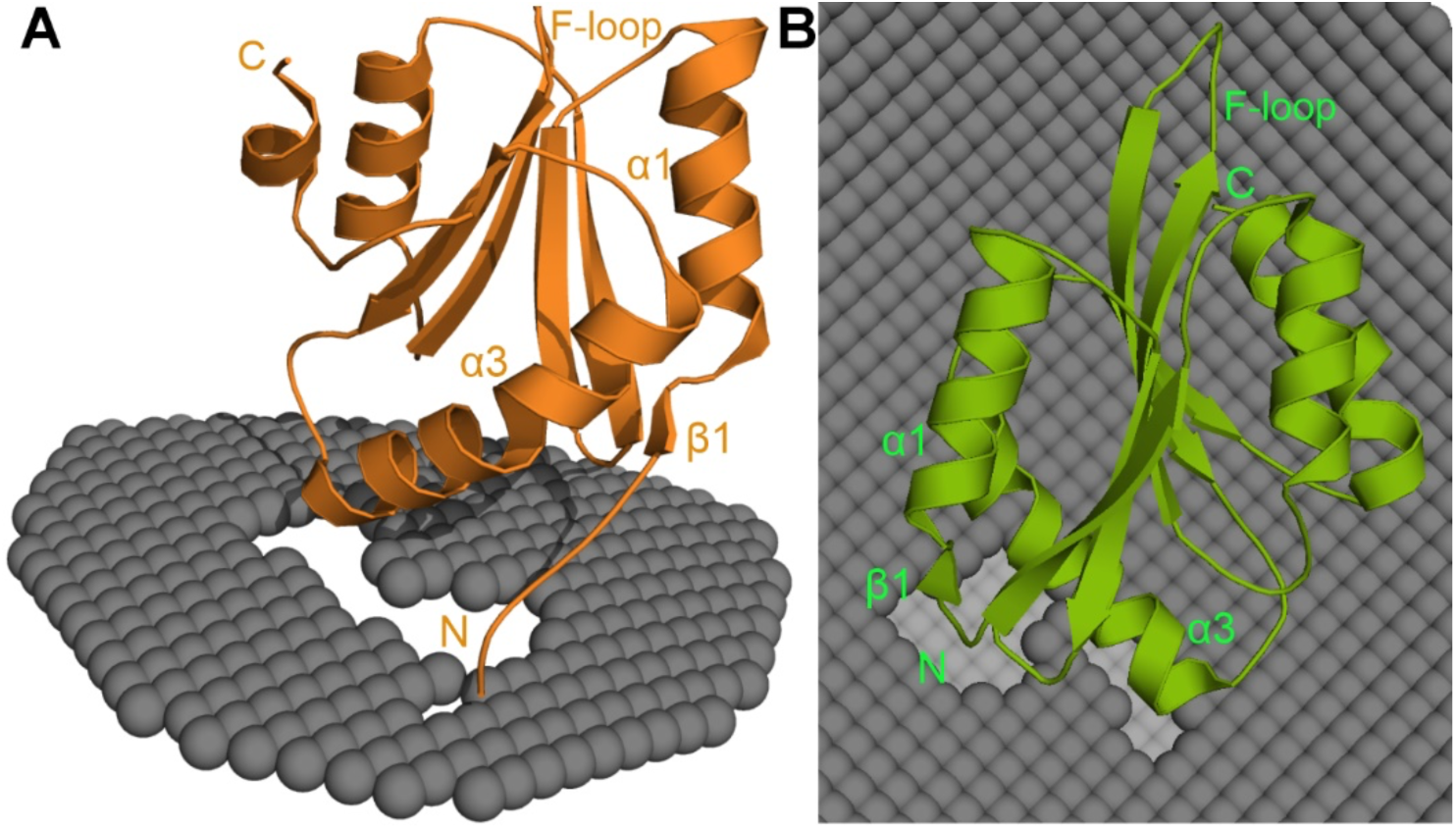
Prediction of membrane binding by PPM. (A) PPM docking of *Pf*ADF1 (A) and *Pb*ADF2 (B) on mammalian plasma membrane. In both models, the N-termini and α3 interact with the membrane.

A recent study on different actin-binding proteins showed that the ADF/cofilins bind 3–6 phosphoinositide head groups and that the interaction is transient (21). The *Pf*ADF1 crystal structure contains four sulphate ions that might mimic phosphoinositide-binding site (28). Interestingly, these sulphates interact with Arg-6, Arg-21, Lys-100, and Lys-101. In this study, mutations to Arg-21, Lys-100, and Lys-101 were found to affect the phosphoinositide binding (Figure 7B, Table 2). Lys-100 of *Pf*ADF1 was found to be crosslinked with SD1 of actin and is thought to be involved in the non-canonical F-actin binding site (43). Thus, phosphoinositide binding to *Pf*ADF1 may inhibit actin binding activity. Here, we could only identify small patches of positively charged residues on the parasite ADFs. In contrast to this, mouse cofilin-1 binds to multiple PI(4,5)P_2_ headgroups simultaneously *via* a large positively charged protein interface that overlaps with the G- and F-actin binding sites (19). This was also observed in the affinity for phosphoinositide. PI(4,5)P_2_ had a K_d_ of approximately 77 µM, which is ∼19-fold lower than that of mouse cofilin-1 which had a K_d_ of 4 µM (19). Vesicles containing POPC/phosphatidylcholine/phosphatidylserine (60:20:20) were used in those experiments, whereas POPC vesicles were used in this study. This might also have contributed to the much lower affinity of *Pf*ADF1.

The N-terminus of the ADF/cofilin proteins is conserved and shown to be important for the F- and G-actin binding. The ADF/cofilins are negatively regulated by phosphorylation (9, 11). The S3E mutation, which mimics phosphorylation at this site, decrease the binding with PI(4,5)P_2_, indicating that phosphorylation might inhibits the phosphoinositide binding (Table 2). In contrast to this study, phosphorylation of chicken cofilin did not affect the affinity to phosphoinositide or the mode of phosphoinositide binding (31). Many studies have shown that the phosphorylation of the N-terminal serine blocks actin interactions and that the S3E mutation made the mutant completely inactive (11, 44, 45). Many cytoskeletal proteins, such as WASP, bind phosphoinositides by basic or aromatic residues (46). As this study identified some basic residues that are involved in the phosphoinositide binding, there might be other residues that may be involved in the phosphoinositide binding, which require further study.

### Experimental procedures

#### Protein expression and purification

Expression constructs for both *Pf*ADF1 and *Pb*ADF2 in the pETNKI-his-SUMO3 vector (NKI Protein Facility, Amsterdam, the Netherlands) with an N-terminal His_6_ tag were obtained from Dr. Moon Chatterjee. Several point mutants (S3A, S3E, K72A, K72Q), double mutants (R86AR88A, R86QR88Q, K100AK101A, K100QK101Q), and a triple mutant (K19AK21AR22A) of *Pf*ADF1 were generated from the *Pf*ADF1 plasmid using polymerase chain reaction with Phusion High-Fidelity DNA polymerase (Thermo Fisher Scientific Inc., Waltham, MA, USA). In each case, the reaction mixture was incubated with DpnI to remove the methylated template. Plasmid DNA was ligated using T4 DNA ligase, followed by transformation to TOP10 competent cells (Invitrogen, Waltham, MA, USA), which were then plated on Luria-Bertani agar plates containing 50 µg/mL kanamycin. Plasmids were isolated from single colonies and screened for the presence of the desired mutation by DNA sequencing at the DNA Sequencing Core Facility at Biocenter Oulu.

Wild type *Pf*ADF1 and the *Pf*ADF1 mutants were transformed into *E. coli* BL21CodonPlus (DE3) RIPL (Agilent, Santa Clara, CA, USA), while *Pb*ADF2 was transformed into *E. coli* Rosetta (DE3) (Novagen, Darmstadt, Germany). Selected transformants were inoculated into Luria-Bertani medium at +37℃ with 50 µg/mL kanamycin and 34 µg/mL chloramphenicol and grown overnight at +37°C. Expression cultures were grown in ZYM-5052 autoinduction medium (47) at +37℃ for 4 h after inoculation with 1% preculture. The cultures were then cooled to +20℃ and incubated for further 36 h for all constructs. The cells were harvested by centrifugation at 5,020 g for 45 min, washed with phosphate-buffered saline and stored at – 20℃. *Pf*ADF1 and *Pb*ADF2 were purified as described before (48). The *Pf*ADF1 mutants were purified following the *Pf*ADF1 purification protocol.

#### Lipid vesicle preparation

Phosphotidylinositol phosphates were purchased from Echelon Biosciences (Salt Lake City, UT, USA). POPC was purchased from Avanti Polar Lipids (Alabester, AL, USA) and DMPC and DMPG from Antrace (Maumee, OH, USA). Lipid stocks of long-carbon-chain phosphoinositide were prepared by dissolving dry lipid in chloroform:methanol:water (20:13:3 v/v), while short-carbon-chain lipids were dissolved in either buffer or water. DMPC:DMPG (1:1) were dissolved in chloroform:methanol (4:1 v/v). Vesicles were prepared by adding deionised water to dried lipids under vigorous shaking. The suspensions were clarified by sonication (Branson 450 Digital Sonifier, Marshall Scientific LLC, Hampton, NH, USA) for 2 min at 10% amplitude with 1 s pulses, forming small unilamellar vesicles (SUV). A mixture of 2 mM POPC:phosphoinositide (90:10), 2 mM POPC, and 5 mg/mL DMPC:DMPG (1:1) were prepared.

#### Vesicle co-sedimentation assay

A vesicle co-sedimentation assay was performed to determine the binding ability of *Plasmodium* ADFs to different phosphoinositide vesicles. 500 µM SUVs were mixed with 8 µM protein in 20 mM 2-[4-(2-hydroxyethyl) piperazin-1-yl] ethanesulfonic acid (HEPES) pH 7.0, 50 mM NaCl in a total volume of 50 µl and incubated for 30 min at room temperature. The samples were then centrifuged at 434,500 g for 1 h at +20°C using an Optima TL-10 benchtop ultracentrifuge (Beckman Coulter, Indianapolis, IN, USA). The supernatant was transferred to a new tube, and the pellet was resuspended in 50 µL of 20 mM HEPES pH 7.0, and 50 mM NaCl. Both supernatant and pellet samples were mixed with 12.5 µl of 5 x SDS-PAGE sample buffer (250 mM Tris pH 6.8, 10% sodium dodecyl sulphate, 50% glycerol, 0.02% Bromophenol Blue and 1.43 M β-mercaptoethanol). The samples were incubated for 5 min at +95°C, and 10 µL of each sample were analyzed on 4%–20% SDS-PAGE gels. The protein bands were visualized using PageBlue staining (Thermo Fisher Scientific Inc.), and the gels were imaged using a ChemiDoc XRS+ system (Bio-Rad, Hercules, CA, USA). Protein band intensities were quantified using the ImageJ software (49). For each supernatant and pellet pair, the total intensity of ADF was set to 100%, and the relative amounts of ADF in the pellets were presented as percentages. The assay was repeated three times.

#### Tryptophan fluorescence assay

Tryptophan fluorescence was used to study the interaction between *Plasmodium* ADFs and different phosphoinositide vesicles. All tryptophan fluorescence spectroscopy experiments were performed in a quartz cuvette with pathlength of 3 mm, using a Fluoromax-4 spectrofluorometer (Horiba Scientific, Kyoto, Japan). The excitation wavelength was fixed to 295 nm, and the emission spectra were collected by averaging ten spectra from 300 to 450 nm at +25℃. An aliquot containing 10 µM ADF was titrated with different concentrations of phosphoinositide vesicles from 0 to 1.5 mM. Spectra of buffer and vesicles alone were subtracted from the ADF alone and ADF-phosphoinositide vesicles sample, respectively. The binding of the ADFs to POPC was also measured as a control. Data from three independent experiments were analyzed using nonlinear regression with ‘one site-specific binding’ model in GraphPad Prism 8 (GraphPad Software, La Jolla, CA, USA) using

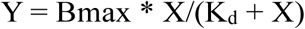

where X is the ligand concentration, Y is the fluorescence intensity, B_max_ is the maximum specific binding, and K_d_ is the equilibrium dissociation constant.

#### Synchrotron radiation circular dichroism

Secondary structure compositions of ADFs in the presence and absence of lipids were determined using SRCD. Lipid vesicles were mixed with protein at a 1:100 protein-to-lipid (P:L) molar ratio prior to the measurement. Protein samples at 0.12 mg/mL were recorded three times in water between 170–280 nm in a Hellma cylindrical absorption cuvette (Suprasil quartz, Hellma GmbH & Co. KG, Müllheim, Germany) with a pathlength of 0.5–1 mm at the AU-CD beamline at the ASTRID2 synchrotron (ISA, Aarhus, Denmark) at +25℃. Buffer spectra were subtracted, and CD units converted to Δε (M^-1^cm^-1^) using CDtoolX (50). Secondary structure deconvolutions were done using BeStSel (30) or DICHROWEB (51).

#### Sequence alignment of *Plasmodium* ADFs with other ADF/cofilin proteins

A multiple sequence alignment of *Plasmodium* ADFs and other ADF/cofilin proteins was generated with ClustalW2 (52) and visualized using ESPript (53). The UniProtKB accession numbers are as follows: *At*ADF1, *A. thaliana* ADF1 (Q39250); *Ac*Act, *Acanthamoeba castellanii* actophorin (P37167); *Sc*Cof, *S. cerevisiae* cofilin (Q03048); *Mm*Cof, *Mus musculus* cofilin-1 (P18760); HsCof, *Homo sapiens* cofilin (P23528); *Pf*ADF1, *P. falciparum* ADF1 (Q8I467); *Pb*ADF2, *P. berghei* ADF2 (Q3YPH0); and *Tg*ADF, *T. gondii* ADF (B9Q2C8).

#### Protein-membrane interaction study

The orientation of the *Plasmodium* ADFs [PDB IDs 2XF1 and 2XFA (28)] on the plasma membrane was estimated using the PPM server version 3.0 (41). A mammalian plasma membrane was used for the calculations, and heteroatoms, water, and detergents were excluded from the calculations. Both *Plasmodium* ADFs were treated as peripheral proteins by the PPM program.

## Data availability

All the plasmids and relevant data used to support the findings of this study are available upon request from the corresponding authors.

## Acknowledgments

We acknowledge the use of the AU-CD beamline on ASTRID2 at ISA (Aarhus, Denmark) for SR- CD measurements. The assistance of Nykola C. Jones and Søren Vrønning Hoffman at the AU-CD beamline is highly valued and appreciated. Access to the facilities and the expertise of the Biocenter Oulu Proteomics and Protein Analysis as well as Structural Biology core facilities, members of Biocenter Finland, are gratefully acknowledged. This work was funded by the Sigrid Jusélius Foundation (I.K.), the Academy of Finland (I.K), the Emil Aaltonen Foundation (I.K), the Jane and Aatos Erkko Foundation (I.K.), and the Norwegian Research Council (I.K.).

## Author contributions

D.L. conceptualization, methodology, formal analysis and validation, writing original draft; I.K. conceptualization, methodology, formal analysis, validation and writing – review and editing, supervision, project administration, and funding acquisition.

## Abbreviations

ADF: actin depolymerizing factor
*Ac*: *Acanthamoeba castellanii*
Act: actophorin
*At*: *Arabdidopsis thaliana*
BeStSel: Beta structure selection
β-ME: beta mercaptoethanol
Cof: cofilin
DMPC: 1,2-dimyristoyl-sn-glycero-3-phosphocholine
DMPG: 1,2-dimyristoyl-sn-glycero-3-phosphoglycerol
*Hs*: *Homo sapiens*
K_d_: dissociation constant
*Mm*: *Mus musculus*
*Pb*: *Plasmodium berghei*
*Pf*: *Plasmodium falciparum*
POPC: 1-palmitoyl-2-oleoly-sn-glycero-3-phosphocholine
PPM: Positioning of Proteins in Membrane
*Sc*: *Saccharomyces cerevisiae*
SDS-PAGE: sodium dodecyl-sulphate polyacrylamide gel electrophoresis
SRCD: synchroton radiation circular dichroism
SUV: small unilamellar vesicle
TCEP: tris(2-carboxyethyl)phosphine
*Tg*: *Toxoplasma gondii*

## References

1. Bamburg, J. R., Minamide, L. S., Wiggan, O., Tahtamouni, L. H., and Kuhn, T. B. (2021) Cofilin and Actin Dynamics: Multiple Modes of Regulation and Their Impacts in Neuronal Development and Degeneration. Cells. 10, 2726

2. Xu, J., Huang, Y., Zhao, J., Wu, L., Qi, Q., Liu, Y., Li, G., Li, J., Liu, H., and Wu, H. (2021) Cofilin: A Promising Protein Implicated in Cancer Metastasis and Apoptosis. Front Cell Dev Biol. 10.3389/fcell.2021.599065

3. Andrianantoandro, E., and Pollard, T. D. (2006) Mechanism of actin filament turnover by severing and nucleation at different concentrations of ADF/cofilin. Mol Cell. 24, 13–23

4. Carlier, M. F., Laurent, V., Santolini, J., Melki, R., Didry, D., Xia, G. X., Hong, Y., Chua, N. H., and Pantaloni, D. (1997) Actin depolymerizing factor (ADF/cofilin) enhances the rate of filament turnover: implication in actin-based motility. J Cell Biol. 136, 1307–1322

5. Wioland, H., Guichard, B., Senju, Y., Myram, S., Lappalainen, P., Jégou, A., and Romet-Lemonne, G. (2017) ADF/cofilin accelerates actin dynamics by severing filaments and promoting their depolymerization at both ends. Curr biol. 27, 1956–1967.e7

6. Nishida, E. (1985) Opposite effects of cofilin and profilin from porcine brain on rate of exchange of actin-bound adenosine 5’-triphosphate. Biochemistry. 24, 1160–1164

7. Moriyama, K., Iida, K., and Yahara, I. (1996) Phosphorylation of Ser-3 of cofilin regulates its essential function on actin. Genes to Cells. 1, 73–86

8. Blanchoin, L., Robinson, R. C., Choe, S., and Pollard, T. D. (2000) Phosphorylation of Acanthamoeba actophorin (ADF/cofilin) blocks interaction with actin without a change in atomic structure. J Mol Biol. 295, 203–211

9. Austin Elam, W., Cao, W., Kang, H., Huehn, A., Hocky, G. M., Prochniewicz, E., Schramm, A. C., Negró, K., Garcia, J., Bonello, T. T., Gunning, P. W., Thomas, D. D., Voth, G. A., Sindelar, C. V, and De La Cruz, E. M. (2017) Phosphomimetic S3D cofilin binds but only weakly severs actin filaments. J Biol Chem. 292, 19565–19579

10. Wioland, H., Jegou, A., and Romet-Lemonne, G. (2019) Quantitative variations with pH of actin depolymerizing factor/cofilin’s multiple actions on actin filaments. Biochemistry. 58, 40–47

11. Pope, B. J., Zierler-Gould, K. M., Kühne, R., Weeds, A. G., and Ball, L. J. (2004) Solution structure of human cofilin: actin binding, pH sensitivity, and relationship to actin-depolymerizing factor. J Biol Chem. 279, 4840–4848

12. Blondin, L., Sapountzi, V., Maciver, S. K., Lagarrigue, E., Benyamin, Y., and Roustan, C. (2002) A structural basis for the pH-dependence of cofilin F-actin interactions. Eur J Biochem. 269, 4194–4201

13. McLaughlin stuart, and Murray Diana (2005) plasma membrane phosphoinsitide organization by protein electrostatics. Nature. 438, 605–611

14. Di Paolo, G., and De Camilli, P. (2006) Phosphoinositides in cell regulation and membrane dynamics. Nature. 443, 651–657

15. Yonezawa, N., Nishidas, E., Iidaq, K., Yaharag, I., and Sakai, H. (1990) Inhibition of the interactions of cofilin, destrin, and deoxyribonuclease I with actin by phosphoinositides. J Biol Chem. 265, 8382–8386

16. Palmgren, S., Ojala, P. J., Wear, M. A., Cooper, J. A., and Lappalainen, P. (2001) Interactions with PIP2, ADP-actin monomers, and capping protein regulate the activity and localization of yeast twinfilin. J Cell Biol. 155, 251–260

17. Barret, C., Roy, C., Montcourrier, P., Mangeat, P., and Niggli, V. (2000) Mutagenesis of the phosphatidylinositol 4,5-bisphosphate (PIP(2)) binding site in the NH(2)- terminal domain of ezrin correlates with its altered cellular distribution. J Cell Biol. 151, 1067–1079

18. Schafer, D. A., Jennings, P. B., and Cooper, J. A. (1996) Dynamics of capping protein and actin assembly in vitro: uncapping barbed ends by polyphosphoinositides. J Cell Biol. 135, 169–179

19. Zhao, H., Hakala, M., and Lappalainen, P. (2010) ADF/cofilin binds phosphoinositides in a multivalent manner to act as a PIP(2)-density sensor. Biophys J. 98, 2327–2336

20. Ojala, P. J., Paavilainen, V., and Lappalainen, P. (2001) Identification of yeast cofilin residues specific for actin monomer and PIP2 binding. Biochemistry. 40, 15562– 15569

21. Senju, Y., Kalimeri, M., Koskela, E. V., Somerharju, P., Zhao, H., Vattulainen, I., and Lappalainen, P. (2017) Mechanistic principles underlying regulation of the actin cytoskeleton by phosphoinositides. Proc Natl Acad Sci U S A. 114, E8977–E8986

22. Senju, Y., and Lappalainen, P. (2019) Regulation of actin dynamics by PI(4,5)P2 in cell migration and endocytosis. Curr Opin Cell Biol. 56, 7–13

23. Sattler, J. M., Ganter, M., Hliscs, M., Matuschewski, K., and Schüler, H. (2011) Actin regulation in the malaria parasite. Eur J Cell Biol. 90, 966–971

24. Schüler, H., and Matuschewski, K. (2006) Regulation of apicomplexan microfilament dynamics by a minimal set of actin-binding proteins. Traffic. 7, 1433–1439

25. Schüler, H., Mueller, A. K., and Matuschewski, K. (2005) A Plasmodium actin-depolymerizing factor that binds exclusively to actin monomers. Mol Biol Cell. 16, 4013–4023

26. Wong, W., Skau, C. T., Marapana, D. S., Hanssen, E., Taylord, N. L., Riglar, D. T., Zuccala, E. S., Angrisano, F., Lewis, H., Catimel, B., Clarke, O. B., Kershaw, N. J., Perugini, M. A., Kovar, D. R., Gulbis, J. M., and Baum, J. (2011) Minimal requirements for actin filament disassembly revealed by structural analysis of malaria parasite actin-depolymerizing factor 1. Proc Natl Acad Sci U S A. 108, 9869–9874

27. Doi, Y., Shinzawa, N., Fukumoto, S., Okano, H., and Kanuka, H. (2010) ADF2 is required for transformation of the ookinete and sporozoite in malaria parasite development. Biochem Biophys Res Commun. 397, 668–672

28. Singh, B. K., Sattler, J. M., Chatterjee, M., Huttu, J., Schuler, H., and Kursula, I. (2011) Crystal structures explain functional differences in the two actin depolymerization factors of the malaria parasite. J Biol Chem. 286, 28256–28264

29. Tawk, L., Chicanne, G., Dubremetz, J. F., Richard, V., Payrastre, B., Vial, H. J., Roy, C., and Wengelnik, K. (2010) Phosphatidylinositol 3-phosphate, an essential lipid in Plasmodium, localizes to the food vacuole membrane and the apicoplast. Eukaryot Cell. 9, 1519–1530

30. Micsonai, A., Wien, F., Bulyáki, É., Kun, J., Moussong, É., Lee, Y. H., Goto, Y., Réfrégiers, M., and Kardos, J. (2018) BeStSel: A web server for accurate protein secondary structure prediction and fold recognition from the circular dichroism spectra. Nucleic Acids Res. 46, W315–W322

31. Gorbatyuk, V. Y., Nosworthy, N. J., Robson, S. A., Bains, N. P. S., Maciejewski, M. W., dos Remedios, C. G., and King, G. F. (2006) Mapping the phosphoinositide-binding site on chick cofilin explains how PIP2 regulates the cofilin-actin interaction. Mol Cell. 24, 511–522

32. Yin, H. L., and Janmey, P. A. (2003) Phosphoinositide regulation of the actin cytoskeleton. Annu Rev Physiol. 65, 761–789

33. Kim, K., McCully, M. E., Bhattacharya, N., Butler, B., Sept, D., and Cooper, J. A. (2007) Structure/function analysis of the interaction of phosphatidylinositol 4,5- bisphosphate with actin-capping protein: Implications for how capping protein binds the actin filament. J Biol Chem. 282, 5871–5879

34. Ojala, P. J., Paavilainen, V., and Lappalainen, P. (2001) Identification of yeast cofilin residues specific for actin monomer and PIP2 binding. Biochemistry. 40, 15562– 15569

35. Yadav, R., Pathak, P. P., Shukla, V. K., Jain, A., Srivastava, S., Tripathi, S., Krishna Pulavarti, S. V. S. R., Mehta, S., David Sibley, L., and Arora, A. (2011) Solution structure and dynamics of ADF from Toxoplasma gondii. J Struct Biol. 176, 97–111

36. Tawk, L., Chicanne, G., Dubremetz, J. F., Richard, V., Payrastre, B., Vial, H. J., Roy, C., and Wengelnik, K. (2010) Phosphatidylinositol 3-phosphate, an essential lipid in Plasmodium, localizes to the food vacuole membrane and the apicoplast. Eukaryot Cell. 9, 1519–1530

37. Ebrahimzadeh, Z., Mukherjee, A., and Richard, D. (2018) A map of the subcellular distribution of phosphoinositides in the erythrocytic cycle of the malaria parasite Plasmodium falciparum. Int J Parasitol. 48, 13–25

38. Wengelnik, K., Daher, W., and Lebrun, M. (2018) Phosphoinositides and their functions in apicomplexan parasites. Int J Parasitol. 48, 493–504

39. Xian, W., Vegners, R., Janmey, P. A., and Braunlin, W. H. (1995) Spectroscopic studies of a phosphoinositide-binding peptide from gelsolin: behavior in solutions of mixed solvent and anionic micelles. Biophys J. 69, 2695–2702

40. Raghunathan, V., Mowery, P., Rozycki, M., Lindberg, U., and Schutt, C. (1992) Structural changes in profilin accompany its binding to phosphatidylinositol 4,5- bisphosphate. FEBS Lett. 297, 46–50

41. Lomize, A. L., Todd, S. C., and Pogozheva, I. D. (2022) Spatial arrangement of proteins in planar and curved membranes by PPM 3.0. Protein Sci. 31, 209–220

42. Prakash, S., Krishna, A., and Sengupta, D. (2023) Cofilin-Membrane Interactions: Electrostatic Effects in Phosphoinositide Lipid Binding. ChemPhysChem. 10.1002/cphc.202200509

43. Wong, W., Webb, A. I., Olshina, M. A., Infusini, G., Tan, Y. H., Hanssen, E., Catimel, B., Suarez, C., Condron, M., Angrisano, F., Nebl, T., Kovar, D. R., and Baum, J. (2014) A mechanism for actin filament severing by malaria parasite actin depolymerizing factor 1 via a low affinity binding interface. J Biol Chem. 289, 4043– 4054

44. Mehta, S., and Sibley, L. D. (2010) Toxoplasma gondii actin depolymerizing factor acts primarily to sequester G-actin. J Biol Chem. 285, 6835–6847

45. Blanchoin, L., and Pollard, T. D. (1998) Interaction of actin monomers with Acanthamoeba Actophorin (ADF/cofilin) and profilin. J Biol Chem. 273, 25106– 25111

46. Tuominen, E. K. J., Holopainen, J. M., Chen, J., Prestwich, G. D., Bachiller, P. R., Kinnunen, P. K. J., and Janmey, P. A. (1999) Fluorescent phosphoinositide derivatives reveal specific binding of gelsolin and other actin regulatory proteins to mixed lipid bilayers. Eur J Biochem. 263, 85–92

47. Studier, F. W. (2005) Protein production by auto-induction in high density shaking cultures. Protein Expr Purif. 41, 207–234

48. Kumpula, E. P., Pires, I., Lasiwa, D., Piirainen, H., Bergmann, U., Vahokoski, J., and Kursula, I. (2017) Apicomplexan actin polymerization depends on nucleation. Sci Rep. 10.1038/s41598-017-11330-w

49. Rueden, C. T., Schindelin, J., Hiner, M. C., DeZonia, B. E., Walter, A. E., Arena, E. T., and Eliceiri, K. W. (2017) ImageJ2: ImageJ for the next generation of scientific image data. BMC Bioinformatics. 18, 1–26

50. Miles, A. J., and Wallace, B. A. (2018) CDtoolX, a downloadable software package for processing and analyses of circular dichroism spectroscopic data. Protein Sci. 27, 1717–1722

51. Miles, A. J., Ramalli, S. G., and Wallace, B. A. (2022) DichroWeb, a website for calculating protein secondary structure from circular dichroism spectroscopic data. Protein Sci. 31, 37–46

52. Larkin, M. A., Blackshields, G., Brown, N. P., Chenna, R., Mcgettigan, P. A., Mcwilliam, H., Valentin, F., Wallace, I. M., Wilm, A., Lopez, R., Thompson, J. D., Gibson, T. J., Higgins, D. G., and Bateman, A. (2007) Clustal W and Clustal X version 2.0. Bioinformatics. 23, 2947–2948

53. Robert, X., and Gouet, P. (2014) Deciphering key features in protein structures with the new ENDscript server. Nucleic Acids Res. 10.1093/nar/gku316

54. Guan, J. Q., Vorobiev, S., Almo, S. C., and Chance, M. R. (2002) Mapping the G-actin binding surface of cofilin using synchrotron protein footprinting. Biochemistry. 41, 5765–5775

55. Zhao, H., Hakala, M., and Lappalainen, P. (2010) ADF/cofilin binds phosphoinositides in a multivalent manner to act as a PIP(2)-density sensor. Biophys J. 98, 2327–2336

