## Supplemental files for "Functional insights into *Plasmodium* actin depolymerizing factor interactions with phosphoinositides"

Materials included:

Figure S1

Figure S2

Table S1

**Figure S1**

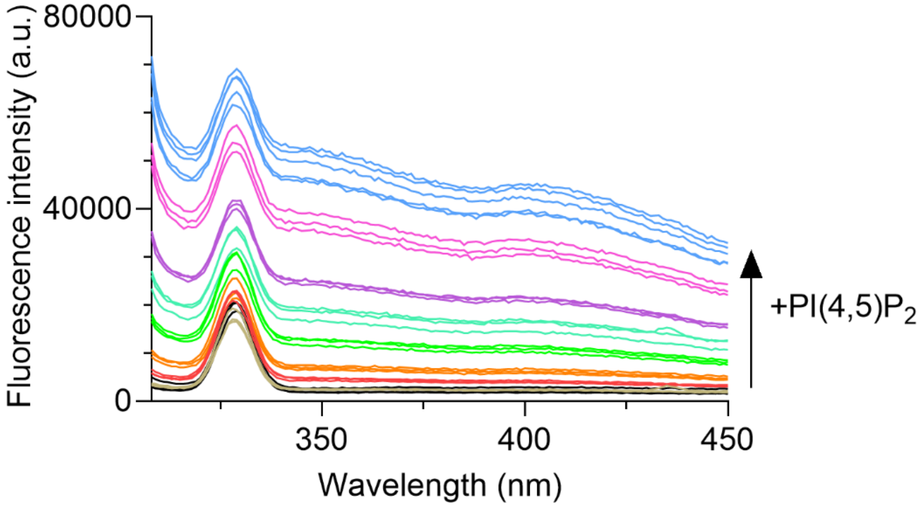


**Figure S1:** Tryptophan emission spectra of PI(4,5)P2 vesicles alone.

**Figure S2**

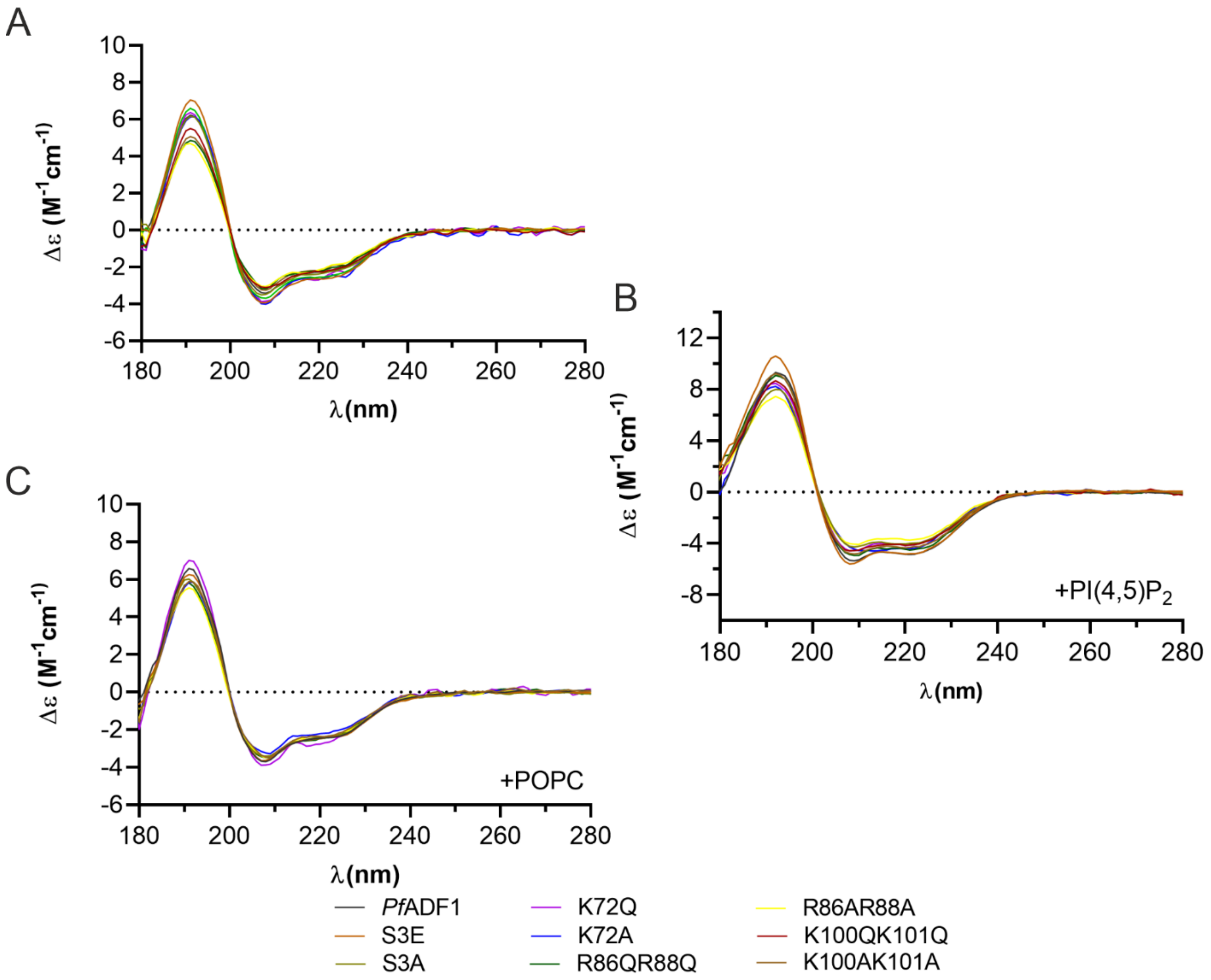


**Figure S2:** SRCD spectra of *Pf*ADF1 mutants in the presence and absence of POPC/PI(4,5)P2. (A) SRCD spectra of the wild-type *Pf*ADF1 and its mutants without PI(4,5)$P_2$. (B) SRCD spectra of *Pf*ADF1 and its mutants with PI(4,5)$P_2$, (C) SRCD spectra of *Pf*ADF1 and its mutants with POPC only.

**Supplementary table S1:** List of the phosphatidylinositol phosphates used.

| Phosphoinositide | Solvent |
|---|---|
| Dipalmitoyl Phosphatidylinositol 3 phosphate | chloroform:methanol:water 20:13:3 |
| Dipalmitoyl Phosphatidylinositol 4 phosphate | chloroform:methanol:water 20:13:3 |
| Dipalmitoyl Phosphatidylinositol 5 phosphate | chloroform:methanol:water 20:13:3 |
| Dipalmitoyl Phosphatidylinositol 4,5-bisphosphate | chloroform:methanol:water 20:13:3 |
| Dipalmitoyl Phosphatidylinositol 3,4-bisphosphate | chloroform:methanol:water 20:13:3 |
| Dipalmitoyl Phosphatidylinositol 3,5-bisphosphate | chloroform:methanol:water 20:13:3 |
| Dipalmitoyl Phosphatidylinositol 3,4,5-trisphosphate | chloroform:methanol:water 20:13:3 |
| Dibutanoyl Phosphatidylinositol 4,5-bisphosphate | water or buffer |
